# Human microbiome-derived peptide affects the development of experimental autoimmune encephalomyelitis via molecular mimicry

**DOI:** 10.1101/2024.07.05.602171

**Authors:** Xin Ma, Jian Zhang, Qianling Jiang, Yong-Xin Li, Guan Yang

## Abstract

**Background:** Gut commensal microbiota has been identified as a potential environmental risk factor for multiple sclerosis (MS), and numerous studies have linked the commensal microorganism with the onset of MS. However, little is known about the mechanisms underlying the gut microbiome and host-immune system interaction.

**Methods:** We employed bioinformatics methodologies to identify human microbial-derived peptides by analyzing their similarity to the MHC II-TCR binding patterns of self-antigens. Subsequently, we conducted a range of *in vitro* and *in vivo* assays to assess the encephalitogenic potential of these microbial-derived peptides.

**Findings:** We analyzed 304,246 human microbiome genomes and 103 metagenomes collected from the MS cohort and identified 731 nonredundant analogs of myelin oligodendrocyte glycoprotein peptide 35-55 (MOG_35-55_). Of note, half of these analogs could bind to MHC II and interact with TCR through structural modeling of the interaction using fine-tuned AlphaFold. Among the 8 selected peptides, the peptide (P3) shows the ability to activate MOG_35-55_-specific CD4^+^ T cells *in vitro.* Furthermore, P3 shows encephalitogenic capacity and has the potential to induce EAE in some animals. Notably, mice immunized with a combination of P3 and MOG_35-55_ develop severe EAE. Additionally, dendritic cells could process and present P3 to MOG-specific CD4^+^ T cells and activate these cells.

**Interpretation:** Our data suggests the potential involvement of a MOG_35-55_-mimic peptide derived from the gut microbiota as a molecular trigger of EAE pathogenesis. Our findings offer direct evidence of how microbes can initiate the development of EAE, suggesting a potential microbiome-based therapeutic target for inhibiting the progression of MS.

**Funding:** National Natural Science Foundation of China (82371350 to GY)

**Research in context:** *Evidence before this study:* On July 31, 2024, we conducted a search on PubMed for articles containing the phrases “gut microbiome and multiple sclerosis” and “gut microbiome and experimental autoimmune encephalomyelitis.” This search yielded a total of 630 and 151 articles, respectively, indicating that the relationship between gut microbiota and the development of MS and EAE is well established. In contrast, our search for “gut microbiome and molecular mimicry and experimental autoimmune encephalomyelitis” revealed only two review papers, highlighting a significant gap in the literature regarding the role of molecular mimicry in connecting gut microbiome dynamics to the development of EAE.

*Added value of this study:* In this study, we employed bioinformatics tools to screen for microbial-derived peptides in the gut that potentially cross-react with autoantigen-specific TCR. Our key findings include: 1) Identification of MOG_35-55_ mimics within the human gut microbiome by employing a combination of TCR-binding footprint screening and prediction model of peptide-MHC II-TCR complexes; 2) Microbial-derived MOG_35-55_ mimics can cross-react with MOG_35-55_-specific CD4^+^ T cells; 3) Among them, peptide 3 predicted from *Akkermansia muciniphila* can induce moderate EAE in mice; 4) Dendritic cells could process and present peptide 3 to MOG-specific CD4^+^ T cells and activate these cells.

*Implications of all the available evidence:* This study suggests the potential involvement of a MOG_35-55_-mimic peptide derived from the gut microbiota as a molecular trigger of EAE pathogenesis. These data may provide a potential microbiome-based therapeutic target for inhibiting the progression of MS.

## Introduction

Multiple sclerosis (MS) is an autoimmune disease characterized by the involvement of autoreactive T cells targeting myelin antigens, leading to the destruction of myelin and damage to axons, resulting in neurologic syndromes and physical disability (1). The global prevalence of MS stands at approximately 2.8 million individuals (2), with an increasing incidence observed in developing countries and among children (3). Although the etiology of MS remains unknown, both genetic and environmental factors have been implicated in its development (1). Accumulating evidence suggests that gut microorganisms play a pathogenic role in MS (4, 5). Research has found notable differences in the composition of gut bacteria between MS patients and healthy individuals (6, 7). Similar alterations in the gut microbiota, including elevated levels of *Akkermansia muciniphila* (*A. muciniphila*), have also been observed in experimental autoimmune encephalomyelitis (EAE), an animal model employed to investigate MS (8). However, how the alterations in the gut microbiota might affect MS/EAE development remains largely unknown.

Molecular mimicry is one of the leading mechanisms in linking the role of microbiota in immune reactivity in autoimmune diseases and cancer (9–13). Reassuringly, it has been reported that microbiota-derived peptides resembling myelin basic protein (MBP) can activate autoreactive T cells in MS (14–16). MBP, along with proteolipid protein and myelin oligodendrocyte glycoprotein (MOG), are myelin antigens targeted by HLA-DR-restricted T cells, leading to MS development (1). Prior studies have demonstrated that hepatitis B virus polymerase can partially mimic MBP, leading to mononuclear cell infiltration in the central nervous systems (CNS) of animals immunized with selected peptide from this viral protein (17). Of note, MOG has been considered a primary target of cellular and humoral immune responses in multiple diseases due to its special extracellular immunoglobulin domain, which is directly accessible to binding partners including antibodies (18–20). Furthermore, the MOG_35-55_ peptide has been widely used to induce EAE. Intrigued by these findings, we hypothesized that microbiome-derived peptides resembling the MOG_35-55_ epitope could trigger the autoimmune response responsible for initiating EAE.

To address this hypothesis, we analyzed human microbiome genomes and metagenome databases collected from the MS cohort. We aimed to identify microbial peptides with MHC II-TCR recognition patterns resembling those of MOG_35-55_, which is presented by the MHC class II I-A^b^ and induces chronic EAE in C57BL/6J mice (21). Subsequently, we identified, synthesized, and tested 8 of these peptides for their ability to activate MOG_35-55_-specific T cells. Our findings revealed that a predicted peptide (P3) derived from the gut commensal organism *A. muciniphila* can activate mouse MOG_35-55_-specific T cells *in vitro*, albeit to a lesser extent compared to the MOG_35-55_ peptide. Furthermore, P3 shows encephalitogenic capacity and has the potential to induce EAE in some animals. Notably, we found that mice immunized with a combination of P3 and MOG_35-55_ developed more severe EAE. Additionally, dendritic cells (DCs) were able to process P3 and present it to CD4^+^ T cells isolated from 2D2 mice expressing a MOG-specific T cell receptor. Our findings suggest that microbial peptide derived from the gut commensal, resembling MOG_35-55_, may mimic the native myelin peptide and potentially contribute to EAE development.

## Material and Methods

### Identify the human MOG_35-55_ peptide analog in the human microbiome genomes and human-associated metagenomic samples

We first collected and curated human microbiome-derived peptide resources, which included human microbiome genomes and human-associated metagenomics samples. In detail, we collected 286,997 human microbiome genomes from Unified Human Gastrointestinal Genome dataset (UHGG, v1.0) (22), and 17249 genomes from CIBIO species-level genome bins (23). Taxonomic assignments were performed for genomes from CIBIO by GTDB-tk (24). In addition, we downloaded 103 metagenomes from MS patients in National Bioscience Database Center (NBDC) Human Database with the accession number of hum0197 (25). The data obtained from the UHGG and NBDC were acquired in accordance with human research ethics guidelines and were approved by the Human Ethics sub-Committee of the City University of Hong Kong (jcc2223ay001). For these metagenomic samples, we applied metagenomic assembly to obtain contigs using MEGAHIT (26) for each sample separately. Taxonomic classification was further conducted for the resulting contigs using CAT (v5.2.3) (27). Taxonomic assignments for genomes or contigs based on the Genome Taxonomy Database (28). Finally, all open reading frames were predicted in each genome or contig using Prodigal-short (29). Human MOG_35-55_ sequence analog searching was conducted by BLASTp with an e-value threshold of 0.001 in a pool of proteins from the above Human microbiome-derived peptide resources.

### The structure-based prediction of peptide-MHC II-TCR binding

Identified analogs were first filtered based on whether they contained the motifs with [YR..F.RV.], [YR..F.R..], [YR..F..V.], [YR….RV], and [Y…F.RV.] are involved in the peptide-MHC-TCR interactions. Next, a special peptide region (∼ 21 residues) containing the above binding pattern was truncated from these filtered analogs. The yielded nonredundant truncated peptides were subjected to further analyses.

The peptide-MHC II-TCR binding prediction was performed in two steps: the peptide-MHC II binding specificity prediction and TCR: peptide-MHC II interactions modeling. Because the advanced peptide-MHC II structure prediction tools often predict the binding structure composed by the binding core of peptides (normally with 9 residues) and receptors (MHC II or TCR) (30–34), these nonredundant truncated peptides were further chopped into a series of consecutive 11-residue windows along peptide with the putative core nine-mer residues and their immediate neighboring residues. The resulting 11-residue windows were subjected to a fine-tuned AlphaFold (35) to predict peptide-MHC II binding with the following parameters: --model_names model_2_ptm_ft --ignore_identities (30). Peptide-MHC II templates (4p23_MH2_H2ABa_H2ABb.pdb,6mng_MH2_H2ABa_H2ABb.pdb,1muj_MH2_H2A Ba_H2ABb.pdb) in the PDB template set were used (30). Mean Inter-chain lowest predicted aligned error (PAE) terms corresponding to the peptide-MHC interactions ≤ 4.34 and peptides’ per-residue confidence score (pLDDT) corresponding to local structural accuracy ≥ 90 were used to classify MHC II binder and non-binder. Next, peptide-MHC II-TCR binding for the predicted binders was modeled by a specialized version of the neural network predictor AlphaFold and TCR–pMHC complex templates-based method, TCRDock with ‘model_2_ptm’ and ‘--benchmark’ parameter set (32). Based on TCRDock’s workflow, three simulations were conducted for each target peptide-MHC II-TCR complex and finally chose the model with the lowest PAE between the TCR and pMHC. Given the hypervariable gene segments in TCR alpha and beta chain largely determine the structure modeling (36), we took TCR:pMHC II structural templates (PDB IDs: 3c60, 3rdt, 3c5z, 6mnn), pairing with four types of mouse TCR, to model the peptide-MHC II-TCR structures. The predicted structures were visualized by PyMoL (Version 2.5.3).

### Mice used in this Study

Six to eight-week-old C57BL/6J (RRID: MGI: 2159769) mice were obtained from the Laboratory Animal Research Unit (LARU) of City University of Hong Kong. The 2D2 TCR transgenic mice C57BL/6-Tg (Tcra2D2, Tcrb2D2) 1Kuch/J (RRID: IMSR_JAX: 006912) were purchased from The Jackson Laboratory and bred in LARU. Mice were housed under a 12-hour light/dark cycle in individually ventilated cages. Food and water were provided ad libitum throughout the study.

### Ethics

All experimental procedures were approved by the Animal Ethics Committee of the City University of Hong Kong (approval number AN-STA-00000493) and performed in accordance with LARU Guidelines.

### Reagents and resources

Reagents and resources are summarized in Supplementary Table 1.

### Active EAE induction

For induction of active EAE, six to eight-week-old mice of both sexes were immunized subcutaneously at two sites on the back flank area with 200 μg of P3 (TTLSFYRPPFLRVRRPFYIIF) or MOG_35–55_ (MEVGWYRSPFSRVVHLYRNGK) peptide (Thermo Fisher, J66557.MCR) emulsified in 200 μl incomplete Freund’s adjuvant (BD bioscience, 263910) supplemented with 2 mg/ml heat-killed *Mycobacterium tuberculosis* (BD bioscience, 231141). Mice then received intraperitoneal 400 ng of pertussis toxin (Calbiochem, 516561) at 0- and 2-day post-immunization (dpi). To check if P3 could aggravate classic EAE, mice were immunized with either the combination of P3 and MOG_35–55_ (200 μg P3 + 50 μg MOG_35–55_), MOG_35-55_ (50 μg), or the combination of a scrambled peptide (KGNRYLHVVRSFPSRYWGVEM) and MOG_35–55_ (200 μg scrambled peptide + 50 μg MOG_35–55_). Priori calculation was not performed, and sample sizes were determined from prior preclinical studies. EAE severity was scored daily in a blinded manner across all groups using the standard scale: 0, no disease; 0.5, tail tip limp; 1, loss of tail tone; 1.5, one leg shows weakness; 2, both legs show weakness with some movement; 2.5, one leg is limp; 3, hind limb paralysis; 3.5, hind limb paralysis and limited movement of waist; 4, hind limb paralysis and forelimb weakness; 4.5, both hind limb and forelimb paralysis; 5, moribund/death (37).

### Antigen recall assay

The peripheral T cell antigen recall response was performed in splenocytes collected from EAE mice at 10 dpi. Mice were anesthetized by 3% isoflurane inhalation and then euthanized by cervical dislocation. Spleens were then collected from the euthanized mice and crushed through a 70-μm cell strainer and centrifuged at 400 × g for 5 min at 4 °C, then incubated in red blood cell lysis buffer (Thermo Fisher, A1049201) for 5 min. After centrifuge, isolated cells were plated at a density of 5×10^5^ cells/well and stimulated with synthesized peptide or MOG_35–55_ (20 μg/ml final concentrations) in RPMI 1640 complete medium (RPMI1640 (Thermo Fisher, 11875093), 10% FBS (Thermo Fisher, A5256801), 2 mM L-glutamine (Thermo Fisher, A2916801), 10 mM Hepes (Thermo Fisher, 15630130), 50 μM 2-mercaptoethanol (Sigma-Aldrich, M6250), 100 IU penicillin (Solarbio, A8180), and 100 μg/mL streptomycin (Solarbio, S8290)), in a total volume of 200 μL per well. Cells were stimulated for 18-hour at 37 °C and 5% CO_2_ atmosphere as previously described (38), then analyzed by flow cytometry for cytokine (IFN-γ, IL-17A) producing cells. To check the antigen recall response of the CNS-infiltrated T cell, spinal cord was collected after euthanizing mice on 17 dpi. Spinal cord tissues were minced and incubated in RPMI 1640 supplemented with 2% FBS, 400 U/ml collagenase IV (Thermo Fisher, 17104019), and 20 μg/ml DNase I (Thermo Fisher, 18047019) for 45 min at 37C°C with agitation. The digested tissues were filtered through a 70-μm filter, and the myelin debris were removed by resuspending the pellet after centrifuge in 40% Percoll (Cytiva, 17089109) and overlaid onto 70% Percoll followed by centrifugation at 860xg for 20 min at 20C°C. Leukocytes at the interface were collected.

### Passive EAE induction

Spleens were collected from mice immunized with MOG_35-55_ at 10 dpi. Splenocytes were restimulated with either MOG_35-55_ or P3 (50 μg/ml) and incubated in RPMI 1640 complete medium at a concentration of 5 × 10^6^ cells/ml with the presence of IL-12 (2 ng/ml) for 72 h at 37 °C. Cells were then collected and resuspended in PBS. A total of 2 × 10^7^ stimulated cells were intraperitoneally injected into C57B/L6J recipients. The clinical scores were graded similarly to active EAE.

### Histopathology

Inflammation and demyelination of spinal cords were assessed on day 17 after immunization. The lumbar enlargement of the spinal cords was isolated from animals and instantly fixed in 4% paraformaldehyde for a minimum of 48 h, then embedded in paraffin and sectioned at 5-μm thickness. Sections were stained with H&E (Solarbio, G1120) or LFB (Abcam, ab150675) to evaluate inflammation and demyelination. The sections were scanned using a NanoZoomer S60 Digital slide scanner (Hamamatsu, C13210-01). The degrees of inflammation (H&E scores) were calculated as 0, normal; The severity of demyelination (LFB scores) was analyzed as 0, normal; 1, myelin sheath injuries in rare areas; 2, a few areas of demyelination; 3, confluent perivascular or subpial demyelination; 4, massive demyelination involving one half of the spinal cord; and 5, extensive demyelination involving the whole spinal cord (39).

### Antigen presentation assays

To obtain stimulator DCs, single cell suspensions from the spleen collected from C57BL/6J mice were prepared by mechanical disruption on a 70-μm cell strainer. Cell suspensions were resuspended in MACS buffer (0.5% BSA (Beyotime, ST025), 2 mM EDTA (Solarbio, E8040) in PBS) after lysing red blood cells. CD11c^+^ cells were then negatively selected according to the manufacturer’s instructions using the mouse Pan Dendritic Cell Isolation Kit (Miltenyitec, 130-100-875). Isolated DCs were pulsed with either PBS, P3, or MOG_35-55_ at 20 μg/ml final concentrations, using 2 x 10^4^ cells in a 96-well plate for 1 h at 37 °C. Cells were then washed extensively and used for co-culture with responder 2D2 CD4^+^ T cells. Responder MOG_35-55_-specific CD4^+^ T cells were isolated from 2D2 TCR transgenic mice spleen by using CD4^+^ T cells Isolation Kit (Miltenyitec, 130-104-454). Sorted CD4^+^ T cells were labeled with Carboxyfluorescein succinimidyl ester (CFSE, Thermo Fisher, C1157) following the user guide. A total of 10^5^ CFSE labeled 2D2 CD4^+^ T cells were then seeded in triplicate in a round bottom 96-well plate with 2 x 10^4^ stimulated DCs (5:1 ratio). DCs and T cells were cultured together for 72-hour at 37 °C and 5% CO_2_ atmosphere, then cells were prepared for flow cytometry to detect T cell proliferation and RNA from these cells was collected for checking *Il17a* and *Ifng* mRNA expression level by RT-qPCR.

### Evaluation of DC activation

To determine the activation state of DCs, isolated DCs were primed with either MOG_35-55_, P3, or scrambled peptide (20 μg/ml) for 1 h at 37 °C, followed by culturing these cells in RPMI 1640 complete medium at a concentration of 2 × 10^5^ cells/ml. The cultured DCs were then harvested at different time points after priming and examined for the activation state by checking the expression levels of CD80, CD86, and MHC II using flow cytometry.

### Gene expression analysis by reverse transcription–qPCR (RT-qPCR)

The total RNA of cells was extracted using RNAiso Plus (Takara, 9108) and reverse-transcribed into cDNA using HiScript III All-in-one RT SuperMix (Vazyme, R333-01) according to the manufacturer’s protocol. qPCR was performed using Taq Pro Universal SYBR qPCR Master Mix (Vazyme, Q712) on a QuantStudioC7 Pro Dx Real-Time PCR System (Thermo Fisher). Primer sequences are listed in Supplementary Table 2 and were synthesized by Tsingke Biotech. The results are shown as relative expression values normalized to *Gapdh*.

### Flow cytometry

Single cell suspensions from the spleen were obtained by mechanical disruption on a 70-μm cell strainer, then incubated in red blood cell lysis buffer for 5 min. For the staining process, dead cells were stained with Propidium Iodide (Sigma-Aldrich, 537060) or Live/Dead Ghost dye (Tonbo Biosciences, 13-0870-T100). For staining of surface markers, cells were incubated in fluorescently labeled antibodies (CD45 (Biolegend, 103132), CD3 (Biolegend, 100235), CD4 (Biolegend, 100437), CD69 (Biolegend, 104513), TCR V_β_11 (Biolegend, 139003), CD11C (Biolegend, 117307), CD80 (Biolegend, 104725), CD86 (Biolegend, 105039), MHC II (Biolegend, 107628)) for 30 min in FACS buffer (PBS containing 2% FBS) at 4 °C. For intracellular staining, cells were washed with 2 mL wash buffer, then fixed and permeabilized by using Cytofix/Cytoperm™ Fixation/Permeabilization Kit (BD Biosciences, 554714) as manufacturer’s instructions. Fixed cells were incubated in fluorescently labeled antibodies (IFN-γ (Biolegend, 505849), IL-17A (eBioscience™, 25-7177-82)) for 30 min in permeabilization buffer at 4 °C. All data were collected on a BD Celesta flow cytometer (BD Biosciences) and analyzed using FlowJo software (v.10.2, BD Biosciences).

### Statistical analysis

Data were analyzed using GraphPad Prism software (Version 9.5.1) and reported as mean ± SEM unless otherwise noted. Gaussian distributions were verified with the Kolmogorov-Smirnov test. The two-tailed unpaired Student *t-test* was used for determining differences between two groups with equal variances, whereas Welch’s t-test was employed for groups with unequal variances. The statistical differences between more than two groups were analyzed using one-way ANOVA with Tukey’s post-test for multiple comparisons. A Mann–Whitney *U* test was used to assess the significance of EAE clinical scores throughout the disease course. For groups with 3 biological replicates, we used the nonparametric Wilcoxon test for two-group comparisons and one-way nonparametric ANOVA for more than two groups’ comparisons. Statistical significance was defined as a *p*-value < 0.05.

## Results

### Well-defined peptide-MHC II-TCR binding motifs guided microbiome-derived epitope discovery from the human microbiome

To uncover the microbiome-derived epitopes resembling the immunodominant MOG_35-55_ peptide, we designed a genome mining workflow involving an analysis of a vast dataset comprising 304,246 human microbiome genomes (23, 40) and 103 metagenomic samples from an MS study cohort (41) (Fig 1A). We first extracted 404,181 MOG_35-55_ analogs (Supplementary Table 3) from these resources, followed by a filtering process that retained 21,557 analogs (3,946 non-redundant sequences) possessing well-defined binding residues at specific sequential spacing (Fig 1B, Supplementary Table 4).

**Figure 1.**
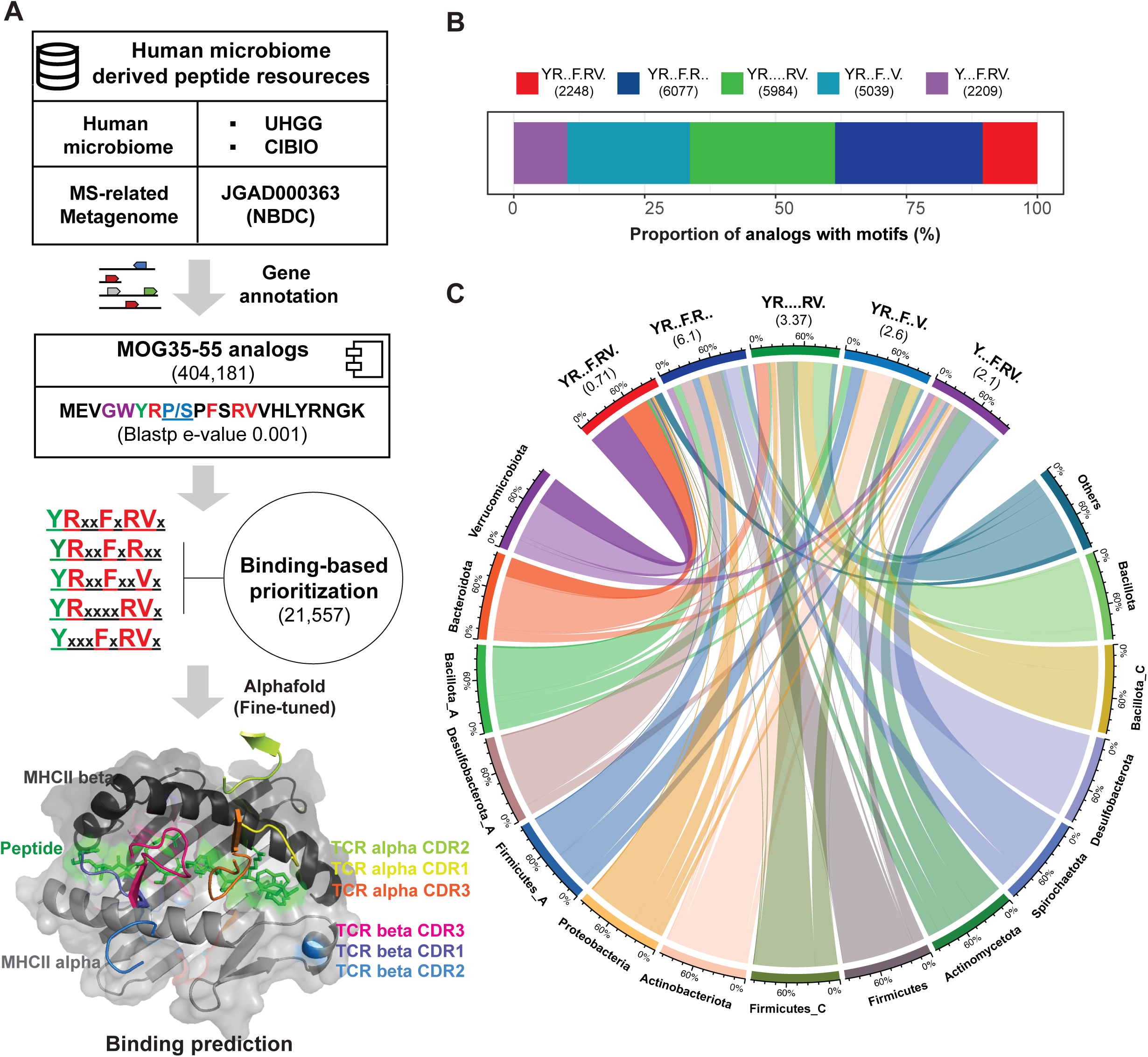
Overview of MOG_35-55_ analogs identified in the human microbiome. (A) The workflow of human microbiome-derived peptides discovery. (B) The proportion of MOG_35-55_ analogs with well-defined binding motifs. (C) Chord diagram showing the taxonomy distribution of prioritized analogs. The values in brackets show the total number of analogs with corresponding binding motifs. per genome in each phylum. The counts of analogs in each phylum were normalized by the genome counts. The scale was the proportion of analogs in each motif (upper) or in each phylum. Of note, the genomes less than 10 counts from *Firmicutes_I*, *Firmicutes_B*, *Bacillota_B*, *Cyanobacteria*, *Desulfobacterota_I*, *Fusobacteriota*, *Pseudomonadota*, *Euryarchaeota*, *Halobacterota*, *Patescibacteria*, and 19 unclassified contigs were merged into Others.

To gain insight into the phylogenetic distribution of MOG_35-55_ analogs derived from the human microbiome, we examined the 21,557 analogs with well-defined binding motifs. Those analogs exhibited a wide distribution across 24 phyla, with the occurrence of a specific motif per genome varying from 0 to 1 within each phylum (Fig 1C). Of note, 10 phyla with fewer than 10 genomes and 19 unclassified bacteria-derived contigs at the phylum level were grouped as “Others”. Interestingly, Verrucomicrobiota demonstrated a higher likelihood of encoding analogs featuring the YRxxFxRVx motif. Besides, these analogs were predominantly encoded by species such as *A. muciniphila* or *A. muciniphila.B*, which are enriched in patients with MS (42).

### Microbiome-derived peptide-MHC II-TCR complexes structure-prediction

Following the general model of antigen presentation (43, 44), we employed a fine-tuned AlphaFold with a binding classifier (30) to discriminate ligands with mouse MHC II molecules (H2-Ab) binding core peptides. Subsequently, the binding core peptides were subjected to modeling the peptide-MHC II-TCR complex by TCRdock (32). First, the performance of fine-tuned AlphaFold and TCRdock was assessed using the MOG_35-55_ peptide as a test case (Fig 2). MOG_35-55_ peptide was further chopped into 10 unique putative binding core peptide candidates (Fig 2A) for subsequent analyses. This was necessary because the current state-of-the-art machine learning-based structure predictors (30, 32–34) for the peptide-MHC II-TCR complex are designed to predict binding between a 9-mer binding core peptide and its receptor (MHC II or TCR). Finally, the binding core peptide of MOG_35-55_, with the specific sequence “WYRPPFSRVVH”, was predicted to be MHC II binders (Fig 2A), and this binding core peptide exhibited reliable interactions with four TCRs that contained different gene segments (Fig 2B). It is noteworthy that this binding core peptide of MOG_35-55_ could interact with mouse MHC II-TCR complex in our pMHC II-TCR model with observed hydrogen bonds (Fig 2C and 2D), which is consistent with the previous studies (21, 45). Inspired by these results, we integrated these two methods to prioritize the microbiome-derived epitopes.

**Figure 2.**
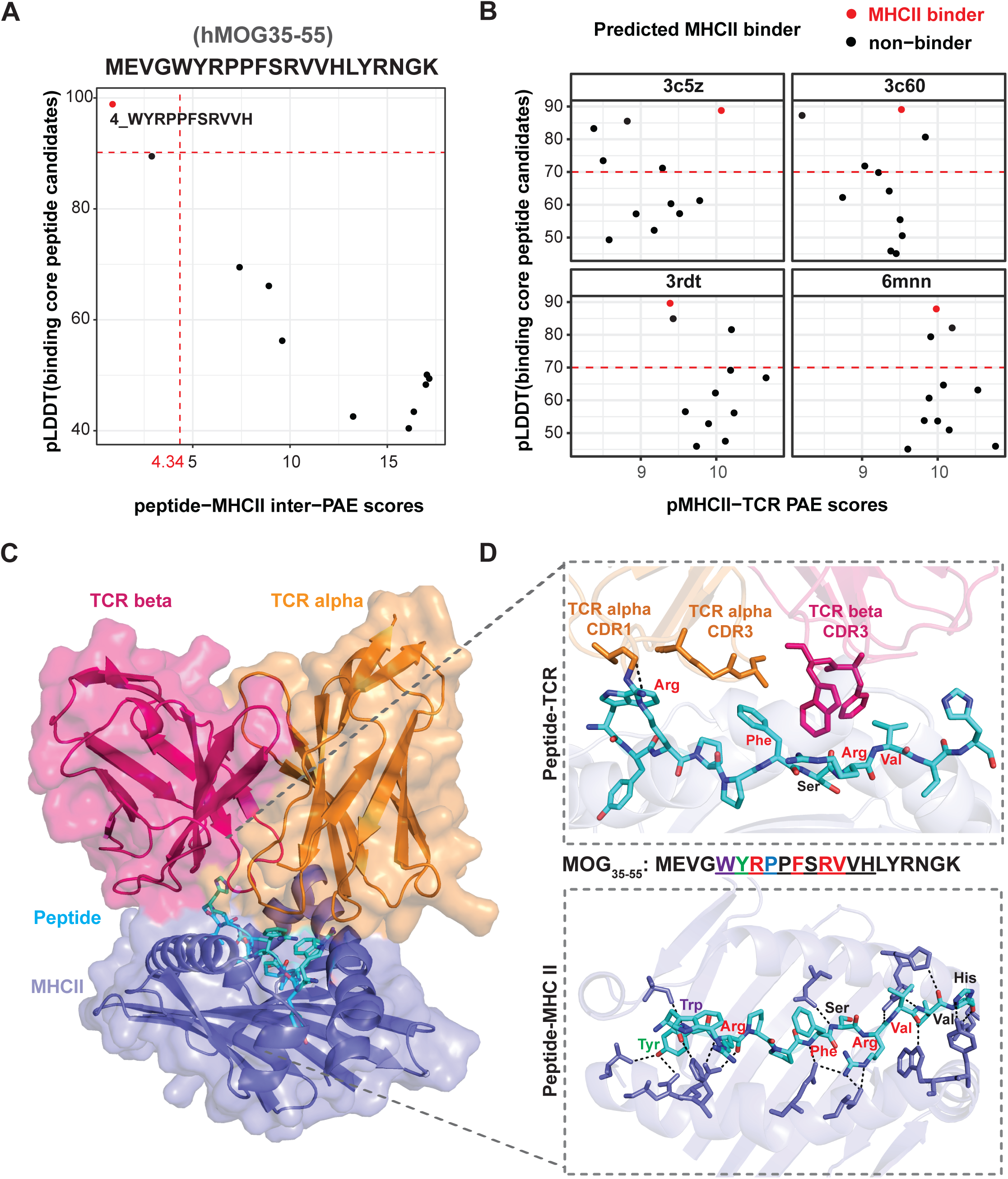
The confidence of hMOG_35-55_ binding core peptide-MHC II and peptide-MHC II-TCR complexes predicted TCRDock. The scatter plots show the peptides’ per-residue confidence score (pLDDT) value of a peptide versus an averaged inter-chain PAE of peptide-MHC-II complexes (A) and peptide-MHC II-TCR complexes (B) for binding core peptide candidates chopped from human MOG_35-55_. The predicted MHC II binding peptides (pLDDT ≥ 90, inter-PAE ≤ 4.34) are highlighted in red. The sequence of binding core peptide candidates is listed on the right side of panel A. The modeling of a peptide with pLDDT≥ 70 is considered as confident. (C) Predicted structure of peptide-MHC II-TCR complex with binding core peptide derived from MOG_35-55_. TCR alleles same from complex template 3rdt (PDB ID). Surface representation is displayed in lighter shades. (D) Close-up view of peptide-TCR interactions (upper) and peptide-MHC II (bottom), colored amino acid residues represent the dominant MHC II-binding residue (green) and major TCR recognition residue (red). Peptide residues and TCR alpha CDR1 and CDR3 (orange) and TCR beta CDR3 (hot pink) residues were shown in sticks, and residues from MHC II interacting with peptide were shown in slate blue. Hydrogen bonds are highlighted in black dashed lines.

To mimic the process of proteolytical degradation of exogenous proteins before their presentation to MHC II in antigen presentation (44), we truncated the corresponding peptide windows (∼ 21 residues) containing the binding motifs along these 3,946 analogs, resulting in 731 nonredundant putative ligand sequences (Fig 3A, Supplementary table 4, 5). These ligands were further chopped into 5,783 unique putative binding core peptide candidates (Supplementary Table 5) for subsequent analyses. Consequently, we identified 493 binding core peptides from 439 ligands as MHC II binders (Fig 3B, Supplementary Table 6). In addition, we successfully modeled interactions between peptide-MHC II and 4 types of TCRs for 97.8% of the MHC II binders using 4 peptide-MHC-TCR complexes templates (Supplementary Fig 1 and Fig 2, Supplementary Table 6). To determine the binding specificity, binding motifs are generated from these binding core peptides and characterized by anchor positions (Fig 3C and 3D). The motifs revealed that Y/F, R, F, R, and V residues are enriched at well-defined anchor positions (loc_1, loc_2, loc_5, loc_7, loc_8) in this study. Surprisingly, several noncanonical binding motifs starting with “L”, “A”, “I”, “S”, “N”, “H”, “Q”, “V”, “T”, “M”, “W”, and others were also observed (Fig 3C). These findings suggest that microbiome-derived epitopes could interact with MHC II and TCRs with diverse binding characteristics.

**Figure 3.**
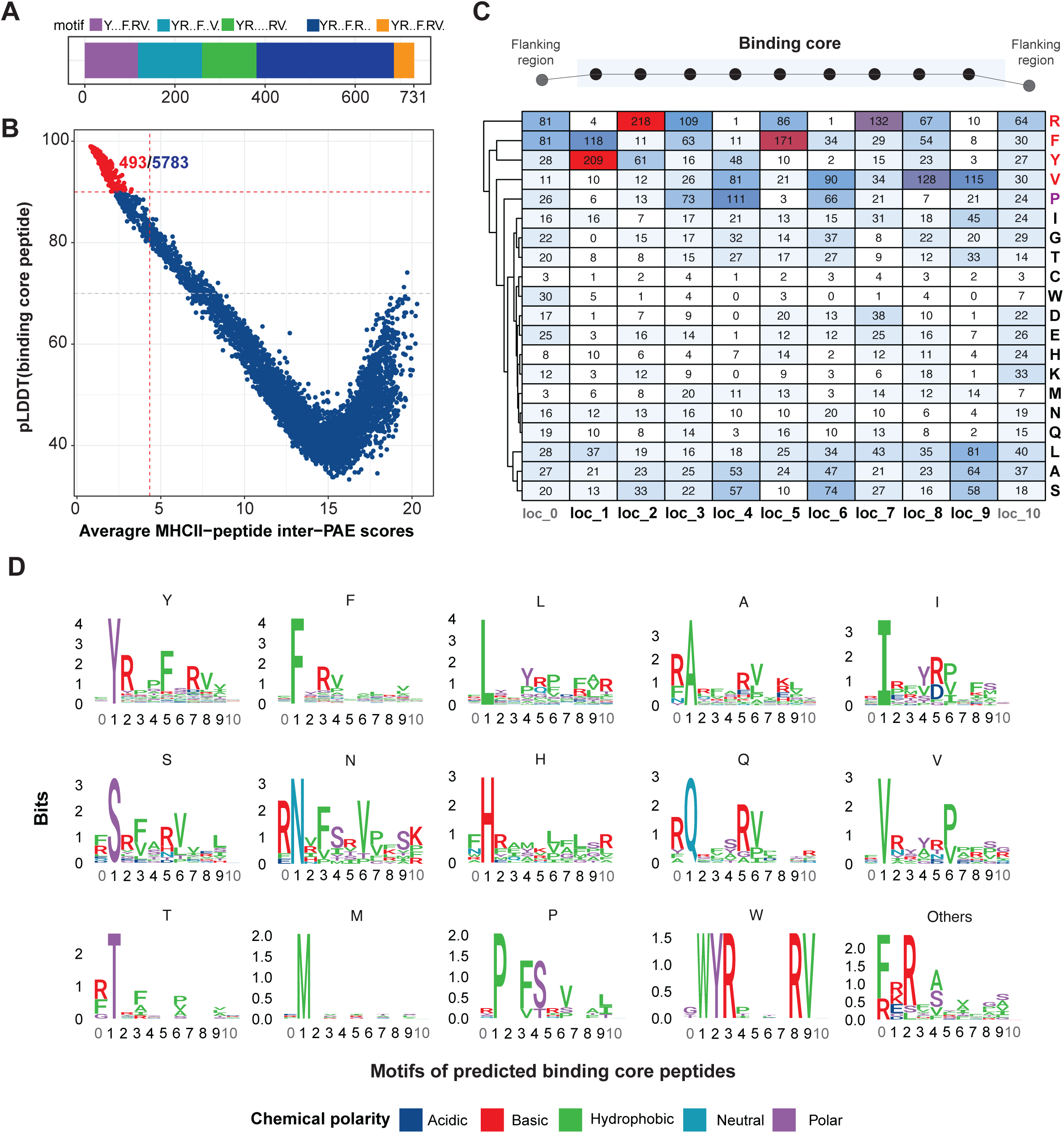
Polymorphism in 9-mer binding core peptides of the predicted peptide-MHC II complex. (A) The stacked bar plot shows the number of nonredundant ligands with 11 amino acids residues truncated from analogs, colored with different motifs. (B) The scatter plot shows the pLDDT value of a binding core peptide (corresponding to local structural accuracy) versus an averaged MHC II-peptide complex’s inter-chain PAE score (corresponding to the peptide-MHC II interactions). The predicted binder is highlighted in red. (C) The frequencies of amino acid residues (rows) at specific positions (columns) among the predicted MHC II binding peptides. The loc_1 ∼ loc_9 represent the binding core region among the binder, and loc_0 and loc_10 represent the flanking region of binders. (D) Representative sequence logos illustrating peptide residue preference of MHC II. Each logo is labeled with the starting amino acid residue of binding core peptides, and the number of binding core peptides used to generate the logos is paired to the corresponding cells in panel [B]. The logos are colored by the chemical polarity of the amino acid residue. In panels C and D, sequence/residue numbers are 0-indexed: they start at 0 and end with N-1, where N is the length of the sequence.

### Microbiome-derived peptide stimulates MOG_35-55_-specific T cell

To investigate the potential modulatory effects of microbiome-derived peptides resembling MOG_35-55_ peptide sequence on the autoimmune response that triggers EAE, a total of 8 peptides (Fig 3 and 4A) were selected from a pool of 439 ligands. These 8 peptides were confidently modeled in pMHC II-TCR interactions (Fig 3), and their selection was based on four specific criteria. Firstly, ligands that contained the “YRxxFxRVx” motif were considered. Secondly, ligands that could be detected in the NBDC database were included (specifically, P1, P2, P4, P5, P6, and P8). Thirdly, ligands collected from species related to MS were chosen (P3 and P5) (42). Finally, ligands containing noncanonical binding motifs starting with “F” and “L” were selected (P1, P3, P7, and P8), as these motifs were found to be most enriched in the pool of 493 binding core peptides. To investigate the potential of microbe-derived peptides to activate MOG_35-55_-specific T cell, we stimulated splenocytes isolated from MOG_35-55_-induced EAE mice with MOG_35-55_ peptide and 8 selected peptides (Fig 4A). Of the tested 8 microbiome-derived peptides, P3 demonstrated the capacity to stimulate the production of IL-17 and IFN-γ by peripheral MOG_35-55_-specific T cells from spleen that were isolated from mice at 10 dpi (Fig 4B-4D, Supplementary fig 3A). Furthermore, P3 also stimulated the production of IFN-γ and IL-17 by spinal cord-infiltrated CD4^+^ T cells during the peak phase of EAE (Fig 4E and 4F). It is noteworthy that P3 was identified as originating from *A. muciniphila*, a gut commensal that is significantly enriched in the gut microbiota of individuals with MS (5).

**Figure 4.**
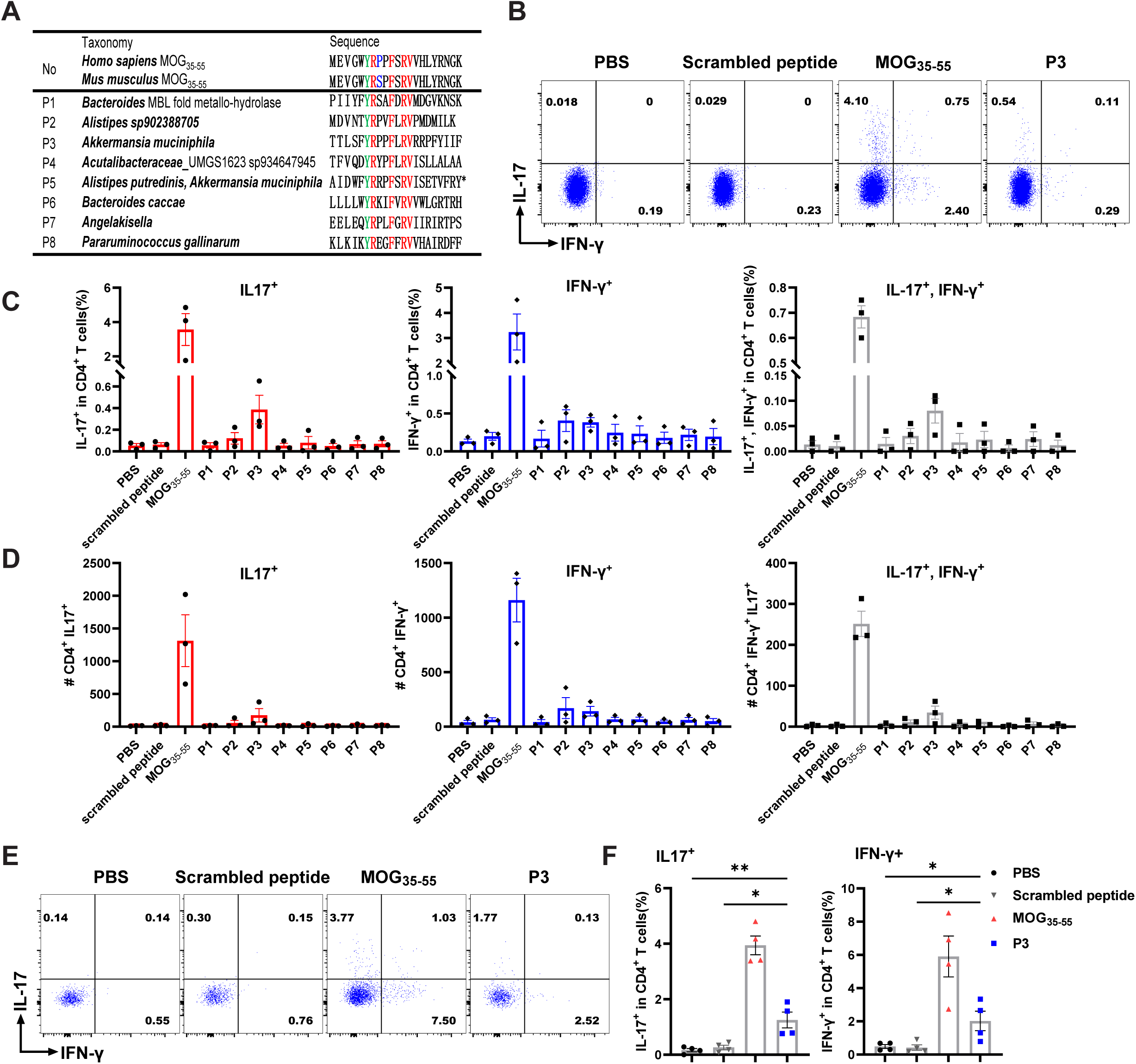
Microbiome-derived peptide can stimulate MOG_35-55_-specific T cells from EAE mice. (A) The origin and sequence of microbial peptides tested in this study. The amino acids highlighted in green are the dominant MHC II-binding residue, while major TCR contact residues are colored in red. The amino acids colored in blue represent different residue between human MOG_35-55_ and mouse MOG_35-55_. The asterisk highlights that P5 is a peptide predicted in both *Alistipes putredinis* and *A. muciniphila*. (B-D) MOG_35-55_–specific CD4^+^ T cells collected from the spleen of EAE mice (n = 3) were treated with each of the 8 selected microbial peptides. PBS and MOG_35-55_ were served as negative control and positive control; Scrambled peptide was set as a peptide control. Flow cytometry was used to determine IFN-γ and IL-17 production. Representative FACS plots (gated on CD3^+^CD4^+^ cells) for the population of IFN-γ- or IL-17-secreted CD4^+^ T cells that were stimulated with mimic peptides or MOG_35-55_ (20 μg/ml final concentration). Only data from P3, which induced the highest cytokine-producing cell population was shown (B), along with the frequency (C) and cell count (D) of each population among CD4^+^ T cells. (E, F) Mice (n = 4) immunized with MOG_35-55_ were sacrificed at 17 dpi. Immune cells from the spinal cord were collected and then stimulated with either MOG_35-55_ (positive control), P3, scrambled peptide, or PBS in a concentration of 20 μg/ml. Representative FACS plots demonstrate the population of activated CD4^+^ T cells (E), along with the frequency (F) of IL-17^+^ and IFN-γ^+^ population among CD4^+^ T cells. Results shown are representative of three independent experiments. Data shown are the average ± SEM. \**p* < 0.05, \*\**p* < 0.01.

### P3 shows encephalitogenic capacity *in vivo*

To further explore the encephalitogenic potential of the P3, we immunized the C57BL/6J mice with P3 (Fig 5A). We found that P3 could induce EAE development in some mice, although both clinical score and incidence were low (disease incidence = 2/12, Fig 5B and 5C). We further stimulated splenocytes from P3-immunized mice sacrificed at 10 dpi with either P3 or MOG_35-55_ for 18-hour (Supplementary fig 4A). Following stimulation, the cells exhibited a robust immune response to P3 and a relatively weaker response to MOG_35-55_ as indicated by the production of IFN-γ (Supplementary fig 4B-4D), suggesting that MOG_35-55_ can cross-react with P3-specific T cells.

**Figure 5.**
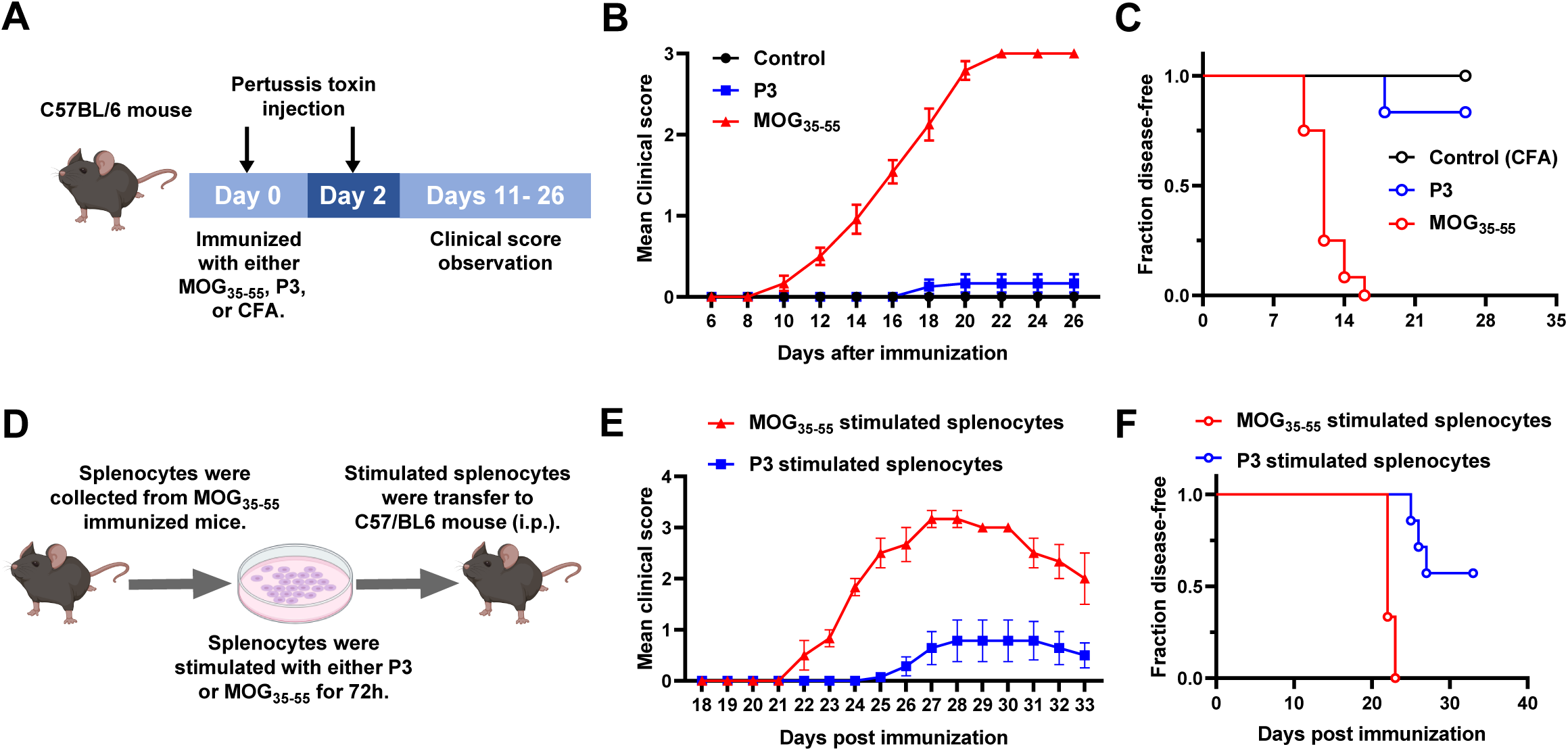
P3 shows encephalitogenic capacity *in vivo*. (A) Experimental design of EAE induction. (B) Mean EAE clinical scores and (C) Kaplan-Meier curve of disease-free survival from mice immunized with either P3, MOG_35-55_, or CFA control. Mice (n = 12) were immunized on day 0 and assessed daily by standard disease scoring for 26 consecutive days. Data shown are pooled from two independent experiments. (D) Experimental outline for adoptive transfer EAE. (E) Mean EAE clinical scores and (F) Kaplan-Meier curve of disease-free survival from mice received either splenocytes stimulated P3 (n = 7) or MOG_35-55_ (n = 3). Data shown are the average ± SEM.

To directly assess the encephalitogenic capacity of P3, we collected splenocytes from MOG_35-55_ immunized mice at 10 dpi and restimulated them with P3. These activated cells were then adoptively transferred to C57BL/6J recipients to induce passive EAE (Fig 5D). As shown in Fig 5E and 5F, recipient mice that received P3-stimulated splenocytes developed EAE with weak symptoms and low incidence. Although the symptoms and incidence were modest, these results suggest that P3 possesses the encephalitogenic capacity.

### P3 in combination with MOG_35-55_ exacerbates EAE development in mice

To further explore the impact of P3 on classic EAE development, C57BL/6J mice were immunized with a combination of P3 and a low dose of MOG_35-55_ (Fig 6A). We found that P3 in combination with MOG_35-55_ could exacerbate EAE development in mice (Fig 6B). To determine the difference of the immune responses in the peripheral between these two groups, we stimulated splenocytes from mice immunized with P3 + MOG_35-55_ or scrambled peptide + MOG_35-55_ with MOG_35-55_. We found that MOG_35-55_ could induce higher levels of IL-17 and IFN-γ in splenocytes collected from mice immunized with P3 + MOG_35-55_ (Fig 6C). At 17 dpi, the spinal cord tissues were collected, and flow cytometry as well as Hematoxylin and Eosin (H&E) or Luxol Fast Blue (LFB) staining were performed to determine the impact of P3 on the severity of inflammation and demyelination. Mice immunized with P3 + MOG_35-55_ displayed more severe inflammation and increased demyelination in the white matter of the spinal cord compared to the other group, with the P3 + MOG_35-55_ group showing significantly higher inflammatory scores (H&E score) and a rising trend in demyelination levels (LFB score) than the Scrambled peptide + MOG_35-55_ group (Fig 6E and 6F). These higher inflammatory scores and demyelination levels in the P3 + MOG_35-55_ group are associated with increased leukocyte infiltration (Fig 6D, Supplementary fig 3B, Supplementary fig 5). We also checked if immunization of P3 could affect the recovery phase of EAE and found that the combination of P3 and MOG_35-55_ showed no significance on the recovery of EAE (Supplementary fig 6A and 6B).

**Figure 6.**
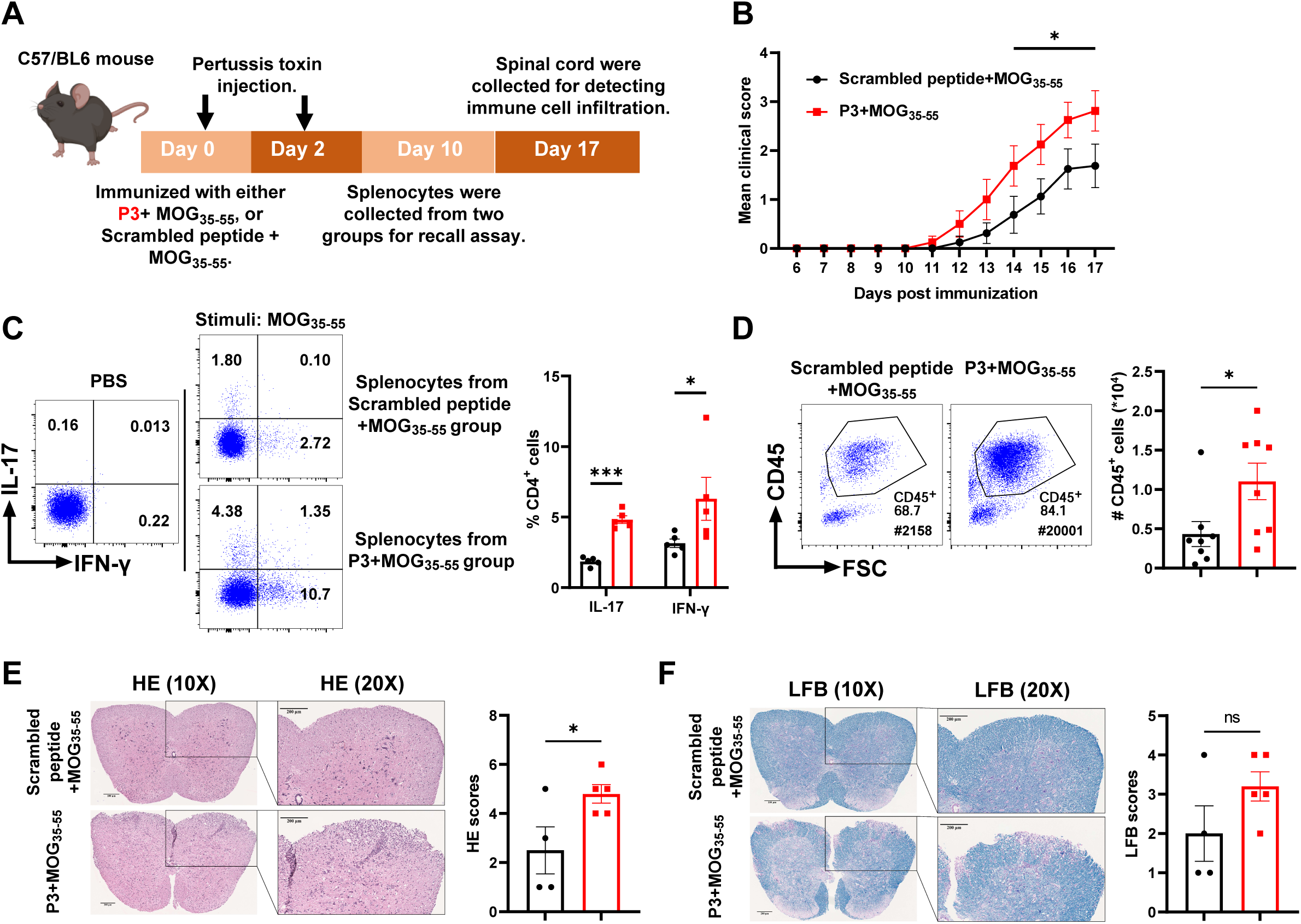
P3 in combination with MOG_35-55_ exacerbates EAE development in mice. (A) The illustration for the experimental design. (B) Mean EAE clinical scores from each group of mice (n = 8). The data shown are representative of two independent experiments. (C) Splenocytes (n = 5) from either P3 + MOG_35-55_ immunized mice or scrambled peptide + MOG_35-55_ immunized mice were stimulated with MOG_35-55_ in a concentration of 20 μg/ml (PBS as a negative stimulation). Representative FACS plots (gated on CD3^+^ and CD4^+^ cells) demonstrate the population of activated CD4^+^ T cells, along with the frequency of IL-17^+^ and INF-γ^+^ population among CD4^+^ T cells. (D) Spinal cord samples were collected at 17 dpi and the leukocyte population within the CNS was assessed by flow cytometry. The cell count of leukocytes was significantly higher in MOG_35-55_ + P3 group than MOG_35-55_ + scrambled peptide group (n = 8). (E, F) Spinal cord tissue sections were obtained at 17 dpi and stained with H&E (E) or LFB (F). The inflammatory level (H&E scores) and demyelination level (LFB scores) were quantified (n=4-5). Data shown are the average ± SEM. \**p < 0.05*, \*\*\**p* < 0.001.

### DCs process microbiome-derived peptide P3 and further activate 2D2 CD4^+^ T cell proliferation

Given that P3 stimulated MOG_35-55_-specific CD4^+^ T cells *in vitro*, we investigated whether DCs are capable of processing and presenting P3 to MOG_35-55_-specific T cells. Our pMHC II-TCR model predicted an interaction between the core 9-mer residues truncated from P3 and the MHC II-TCR complex (Fig 7A and 7B). Hydrogen bonds were observed inside the recognition model similar to the docking result from MHCII-MOG_35-55_-TCR complex (Fig 2C and 2D), leading us to hypothesize that this peptide could be presented by DCs. To test this hypothesis, we performed an antigen presentation assay using DCs isolated from the spleens of C57BL/6J mice. The DCs were primed with either PBS, P3, or MOG_35-55_, and then co-cultured with sorted 2D2 CD4^+^ T cells. The activation status of CD4^+^ T cells was later assessed by quantifying the expression of the early T cell activation marker CD69. Furthermore, the proliferation of CD4^+^ T cells was evaluated by monitoring the changes in the population of CFSE-labeled cells. The results demonstrated that P3 could be presented by DCs and subsequently activate the proliferation of 2D2 CD4^+^ T cells (Fig 7C, Supplementary fig 3C). However, P3 induces a weaker CD4^+^ T cell activation and proliferation than MOG_35-55_ (Fig 7D), which may explain the lower incidence and clinical score observed in P3-induced EAE (Fig 5B and 5C). Furthermore, we noticed that co-culturing P3 with DCs and 2D2 CD4^+^ T cells can enhance the expression of *Il17* and *Ifng* by 2D2 CD4^+^ T cells (Fig 7E). Additionally, we found that P3 could activate DCs by stimulating the expression of CD80 and MHC II (Fig 7F).

**Figure 7.**
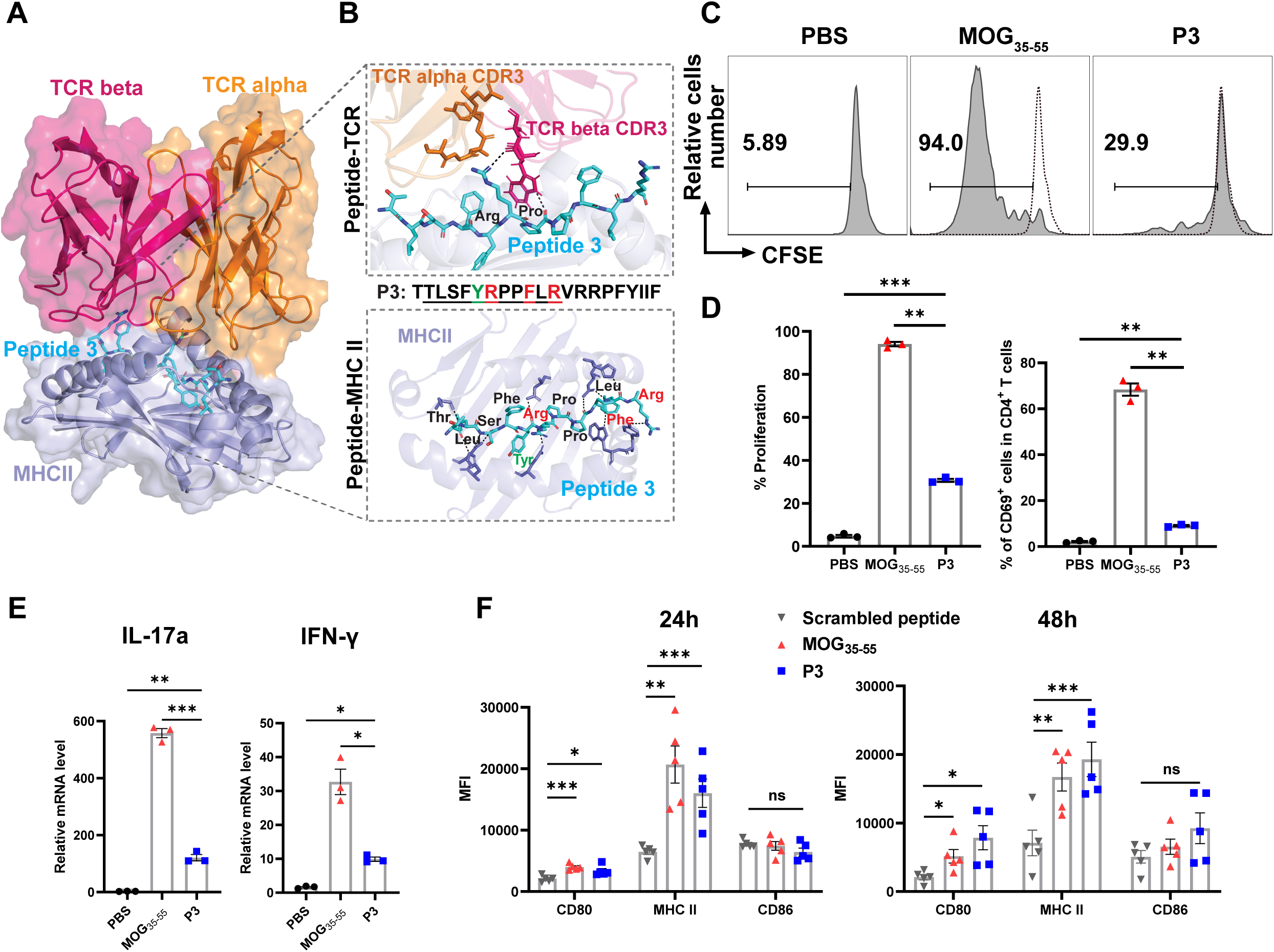
P3 stimulates 2D2 CD4^+^ T cell proliferation through DCs presentation *in vitro*. (A) Predicted structure of peptide-MHC II-TCR complex with binding core peptide from P3. TCR alleles same from complex template 3c60 (PDB ID). Surface representation is displayed in lighter shades. (B) Close-up view of peptide-TCR interactions (upper) and peptide-MHC II (bottom) in panel D, colored amino acid residues represent the dominant MHC II-binding residue (green) and major TCR recognition residue (red). Peptide residues and TCR alpha CDR3 (orange) and TCR beta CDR3 (hot pink) residues were shown in sticks, and residues from MHC II interacting with peptide were shown in slate blue. Hydrogen bonds are highlighted in black dashed lines. (C, D) The histogram FACS plots (C) and quantification (D) of CFSE labeled 2D2 CD4^+^ T cells proliferation (n = 3), as well as the frequency of CD69^+^ cells within the CD4^+^ T cells population, following co-culture with DCs pulsed with PBS, P3, or MOG_35-55_. Dash line in histogram represents the proliferation level of PBS control. (E) The relative mRNA expression of *Il17a* and *Ifng* genes was determined by RT-qPCR. The mean expression levels are shown (nC=C3). (F) DCs were isolated from C57BL/6J mice (n = 5) and stimulated with either MOG_35-55_, P3, or scrambled peptide in a concentration of 20 μg/ml for 24 h and 48 h. The activation of DCs was assessed by measuring MFI of CD80, CD86, and MHC II. Results are representative of two independent experiments. Data shown are the average ± SEM. \**p < 0.05*, \*\**p* < 0.01, \*\*\**p* < 0.001.

## Discussion

The gut microbiota has been implicated in the development of autoimmune diseases. Investigations of MS patients and the EAE mouse model have revealed that the gut microbiome is a crucial factor in disease progression and severity (4, 5). Alterations in gut bacterial composition, including reductions in butyrate-producing bacteria and increases in *Akkermansia* and *Clostridia* populations have been consistently observed in MS patients (5, 7). Moreover, transfer of microbiota from MS patients to animal models has been shown to increase disease incidence, suggesting that alterations in the microbiome may actively contribute to the development of MS (46). Germ-free and antibiotic-treated mice have shown resistance to EAE, further supporting the necessity of the microbiome for disease induction (47, 48).

Despite significant advances in understanding the role of the gut microbiota in MS pathogenesis, the underlying mechanisms leading to disease development remain poorly understood. Continuous sampling of gut microbial antigens by innate immune cells enhances the probability of presenting specific antigens to antigen-specific T cells. The presentation of these antigens is influenced by polymorphisms in MHC genes, suggesting a potential role for host-commensal microorganisms cross-reactivity in the development and persistence of autoimmunity among genetically predisposed individuals. Molecular mimicry has been proposed as a possible mechanism by which microorganisms can induce autoimmune diseases (15). According to this hypothesis, T cells that respond to an epitope derived from an infectious agent may cross-react with a self-antigen that shares sequence or structural similarities with the original microbial peptide (49). Evidence supporting this hypothesis has been observed in numerous autoimmune diseases, such as antiphospholipid syndrome and Type 1 Diabetes (50, 51). Given the diverse range of microorganisms in the gut, peptides derived from gut flora could potentially trigger the development of MS. The activation of autoreactive T cells is a critical component of the early autoimmune response in individuals who ultimately develop MS. While multiple potential antigens exist, MOG is a major EAE antigen in H-2b mice and a possible autoantigen in MS (21). The binding pattern of the MOG-MHC II-TCR complex has been characterized, revealing that tyrosine (Y) in position 40 of the MOG_35-55_ peptide is the dominant MHC II-binding residue, occupying the p1 pocket. Meanwhile, arginine (R) 41, phenylalanine (F) 44, R46, and valine (V) 47 are the major amino acids involved in TCR recognition (21). Research suggests that differences in the amino acid at position 42 between human MOG and mouse MOG may be associated with inducing B cell-dependent EAE (52). Further studies have shown that human MOG fails to induce EAE in mice with B cell antigen-presenting function deficiency (53). These findings collectively underscore the pivotal role of B cells as antigen-presenting cells in developing EAE. However, our current study aims to elucidate how auto-antigen mimic peptides derived from gut commensals stimulate auto-reactive T cells rather than focusing on the specific cell type responsible for presenting these mimic peptides as antigens. Therefore, we excluded this amino acid from our screening motif. Research suggests that low-affinity hotspot mimicry, which involves a subset of crucial residues within the TCR-binding footprint, rather than high-affinity structural mimicry, maybe a more common contributor to the initiation of autoimmune diseases (14, 21). In our study, we focused on identifying MOG_35-55_ mimics based on the similarity of essential residues within the TCR-binding footprint, enabling a broader screening of potential target peptides in the gut microbiome that could induce MS.

Various immune cells play crucial roles in the pathophysiology of MS and EAE. CD4^+^ T cells are central in coordinating the adaptive immune response, releasing cytokines and chemokines that activate myelin-specific CD4^+^ T cells and promote their infiltration into the brain (54). Both IFN-γ-producing Th1 and IL-17-producing Th17 cells can induce EAE by activating CNS macrophages and microglia, leading to local inflammation and tissue damage (55). Our use of the MHC II-peptide-TCR recognition pattern for screening mimic analogs is driven by the pivotal role of CD4^+^ T cells in EAE pathogenesis. Besides CD4^+^ T cells, CD8^+^ T cells also contribute significantly to MS development, as shown by disease induction through the transfer of myelin antigen-specific CD8^+^ T cells in EAE (56). B cells also participate in MS pathogenesis by presenting antigenic peptides to T cells (53), producing cytokines and contributing to the differentiation of lymphocyte subpopulations. Given the diverse functions of B cells, it is possible that gut microbiota mimics could also impact certain B cell-associated autoimmune diseases through the recognition by B cells. Additionally, recent research implicates neutrophils in MS pathogenesis, with their depletion before disease onset ameliorated symptoms (57). Neutrophils recruited to the brain by CD4^+^ T cells can exacerbate parenchymal inflammation. Despite the contributions of various immune cells in MS, CD4^+^ T cells remain central to its pathogenesis. Our study suggests that the exogenous antigens from the gut may activate autoreactive CD4^+^ T cells through molecular mimicry.

By leveraging the expanding genome databases for human gut microbes, we identified 8 microbial peptides that exhibit significant analogy in their antigen-MHC II-TCR binding patterns with MOG_35-55_ peptide. Our findings indicated that microbiome-derived P3 can activate CD4^+^ T cells specific to MOG_35-55_ peptide in mice. During the screening process, peptides like P2 and P4 exhibited only a single amino acid difference from the core sequence of P3, yet did not demonstrate encephalitogenic capacity like P3. This discrepancy may stem from the potential impact of individual amino acid variations on the spatial structure affecting the binding of peptides to MHC II and TCR (58). This observation suggests that in future studies, it may be necessary to evaluate the influence of single amino acid variances on the spatial structure affecting the binding of MHC II-peptide-TCR complex. Interestingly, our bioinformatic analysis revealed that P3 was derived from *A. muciniphila*, suggesting a potential impact of *A. muciniphila* on EAE development. Previous studies have explored the link between *A. muciniphila* and EAE or MS, such as the findings of Cekanaviciute *et al*., who observed that *A. muciniphila* promotes Th1 differentiation in human peripheral blood mononuclear cells (5). Moreover, previous studies have demonstrated that a peptide from *A. muciniphila* shares a similar sequence with an MS-associated self-antigen, guanosine diphosphate (GDP)–L-fucose synthase. This shared peptide sequence has the ability to stimulate the proliferation of CD4^+^ cerebrospinal fluid-infiltrating T cells from MS patients (16). In this study, we optimized the process of screening self-antigen mimics from gut microbiota by integrating the binding affinity to MHC II and TCR into the selection criteria, thereby enhancing the accuracy of identifying self-antigen analogs in the gut microbiome. Furthermore, our findings suggest that the enrichment of *A. muciniphila* in the gut microbiota of EAE mice and MS patients may have pathological implications as this bacterium has the capability of generating MOG_35-55_ analogs.

*A. muciniphila*, a next-generation probiotic, has shown positive effects on several diseases including obesity and diabetes (59). However, recent studies have revealed an enrichment of *A. muciniphila* in the gut microbiome of MS patients. Furthermore, transplantation of fecal samples from MS patients into the intestines of EAE mice leads to more severe EAE symptoms (5), indicating the need to assess the potential beneficial or harmful effects of *A. muciniphila* from multiple perspectives. Although multiple strains of *A. muciniphila* have been identified (60), it remains unclear whether the observed effects in different disease contexts are attributable to strain differences. The predominant use of metagenomics and 16S sequencing in numerous studies exploring the relationship between various diseases and *A. muciniphila* has impeded the identification of specific target strains, as these methods primarily facilitate gene annotation at the species level, presenting challenges in strain-level identification. Our study suggests that *A. muciniphila* may contribute to the development of EAE due to its capacity to generate MOG_35-55_-resembling peptides. However, more evidence is needed to determine the target strain that generates MOG_35-55_ analogs. Future studies will also focus on elucidating the role of synergistic or antagonistic effect among different microbial populations in autoimmune diseases. This study highlights the necessity for further research on the correlation between gut microbiota and autoimmune diseases in the growing trend of widespread probiotic consumption.

In summary, our findings prove a mechanism of molecular mimicry in which a particular sequence within commensal gut microbe can imitate crucial residues in the MOG-MHC II-TCR binding footprint, consequently initiating or modifying the immune response associated with EAE development. While our study focused only on MOG_35-55_ mimics, other myelin antigens, such as MBP and proteolipid protein, may also serve as targets for autoreactive CD4^+^ T cells (61). Given the enormous number of microbial peptides generated by the gut microbiota, it is possible that other microbial peptides may exist and can mimic these antigens and activate related autoantigen-reactive T cells. Meanwhile, it’s important to note that the induction of autoimmune diseases through molecular mimicry is complicated and a single factor acting alone is unlikely to trigger the disease. Instead, accompanying host genetic susceptibility is considered to be linked with an abnormal immune response triggered by one or more microbial antigens or molecules. Nonetheless, this discovery may provide a therapeutic target and an opportunity to impede the progression of MS.

Our data suggests the potential involvement of a MOG_35-55_-mimic peptide derived from the gut microbiota as a molecular trigger of EAE pathogenesis. However, the limitations of this work must be acknowledged. First, while metagenomic approaches are valuable, they have inherent limitations in identifying specific bacteria strains. The prevalent focus on gene annotation at the species level in metagenomics and 16S sequencing restricts strain-level identification (62). Although marker-based computational methods provide strain-level sensitivity (63), they often operate under a single dominant strain per species, thereby overlooking the complex mixture of closely related strains within human-associated microbiota (64). Our methodology utilizing metagenomic data cannot identify the specific *A. muciniphila* strain capable of producing MOG_35-55_ analogs. These challenges underscore a limitation in our ability to identify the precise *A. muciniphila* strain of interest.

Collectively, we have investigated the role of molecular mimicry in the link between microbial flora and EAE development. We have identified MOG_35-55_ mimics derived from gut commensal cross-react with MOG_35-55_-specific CD4^+^ T cells and induce EAE in some mice. The combination immunization of this mimic and MOG_35-55_ significantly exacerbates the development of EAE in mice. Notably, antigen-presenting assays confirmed that DCs can process and present this gut commensal-derived peptide to MOG_35-55_-specific CD4^+^ T cells. Our findings offer direct evidence of how microbes can initiate the development of EAE, suggesting a potential microbiome-based therapeutic target for inhibiting the progression of MS.

## Contributors

Conceptualization: GY, YXL; Methodology: XM, JZ, QLJ; Investigation: XM, JZ; Visualization: XM, JZ; Supervision: GY, YXL; Writing: XM, JZ, YXL, GY. XM and JZ have directly accessed and verified the underlying data reported in the manuscript. GY and YXL are responsible for the decision to submit the manuscript.

## Data Sharing Statement

All study data are included in the article and/or Supplementary files.

## Acknowledgments

This work is supported by the National Natural Science Foundation of China (82371350). The funding source has no role in the writing of the manuscript or the decision to submit it for publication.

