## Supplementary figures for "Human microbiome-derived peptide affects the development of experimental autoimmune encephalomyelitis via molecular mimicry"

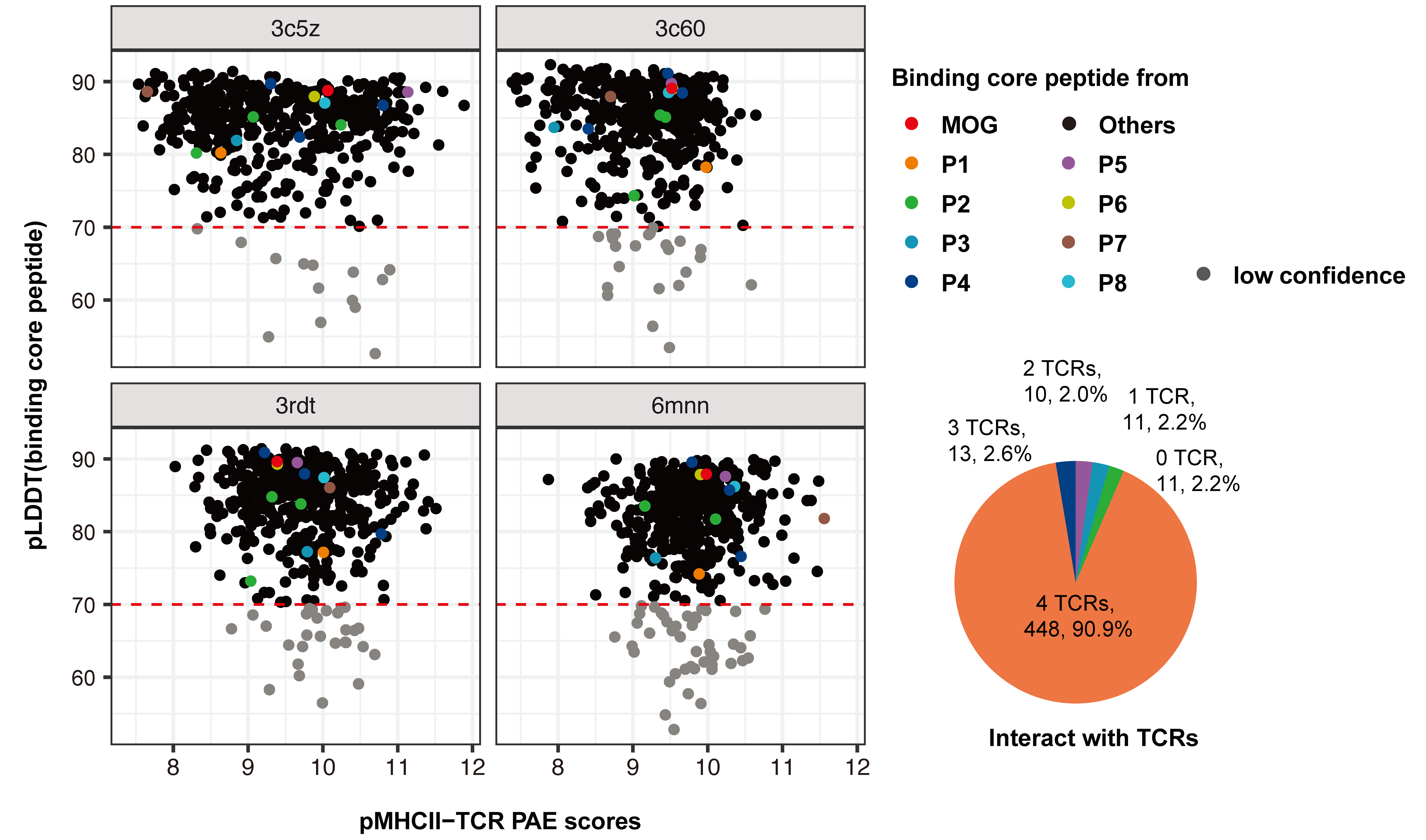

Supplementary figure 1. The confidence of microbiome-derived ligands’ binding core peptide-MHC II interact with 4 types of TCR predicted TCRDock. The scatter plots show the pLDDT value of a peptide versus an averaged inter-chain PAE of peptide-MHCII-TCR complexes. Two binding core peptides of MOG highlighted in green and orange, were shown as controls. Microbiome-derived peptides were colored in red, grey, and grey based on the modeling confidence. The complex with a binding core peptides’ pLDDT ≥ 70 and pMHCII-TCR PAE ≤ 8 was considered confident, and the complex with a binding core peptides’ pLDDT ≥ 70 and pMHCII-TCR PAE > 8 was considered as low confidence. The complex with a binding core peptides’ pLDDT < 70 was considered very low confidence. The pie chart shows a binding core peptide could interact with how many TCRs with confidence or low confidence: 91% of peptides could interact with 4 specific TCRs.

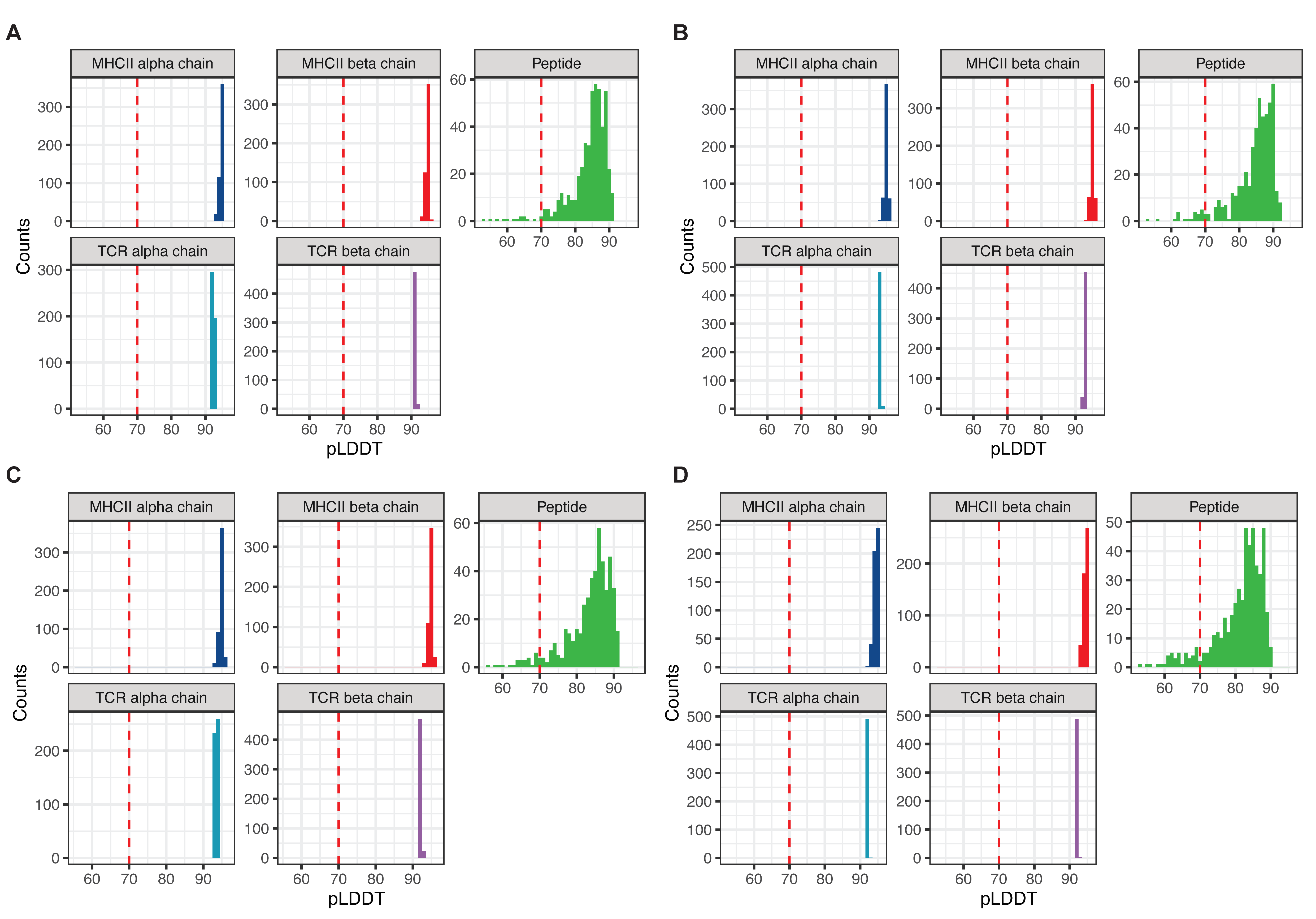

**Supplementary figure 2.** Per-residue confidence scores (pLDDT) of each chain for peptide-MHC-TCR complexes using templates with PDB id: 3c5z (A), 3c60 (B), 3rdt (C), 6mnn (D).

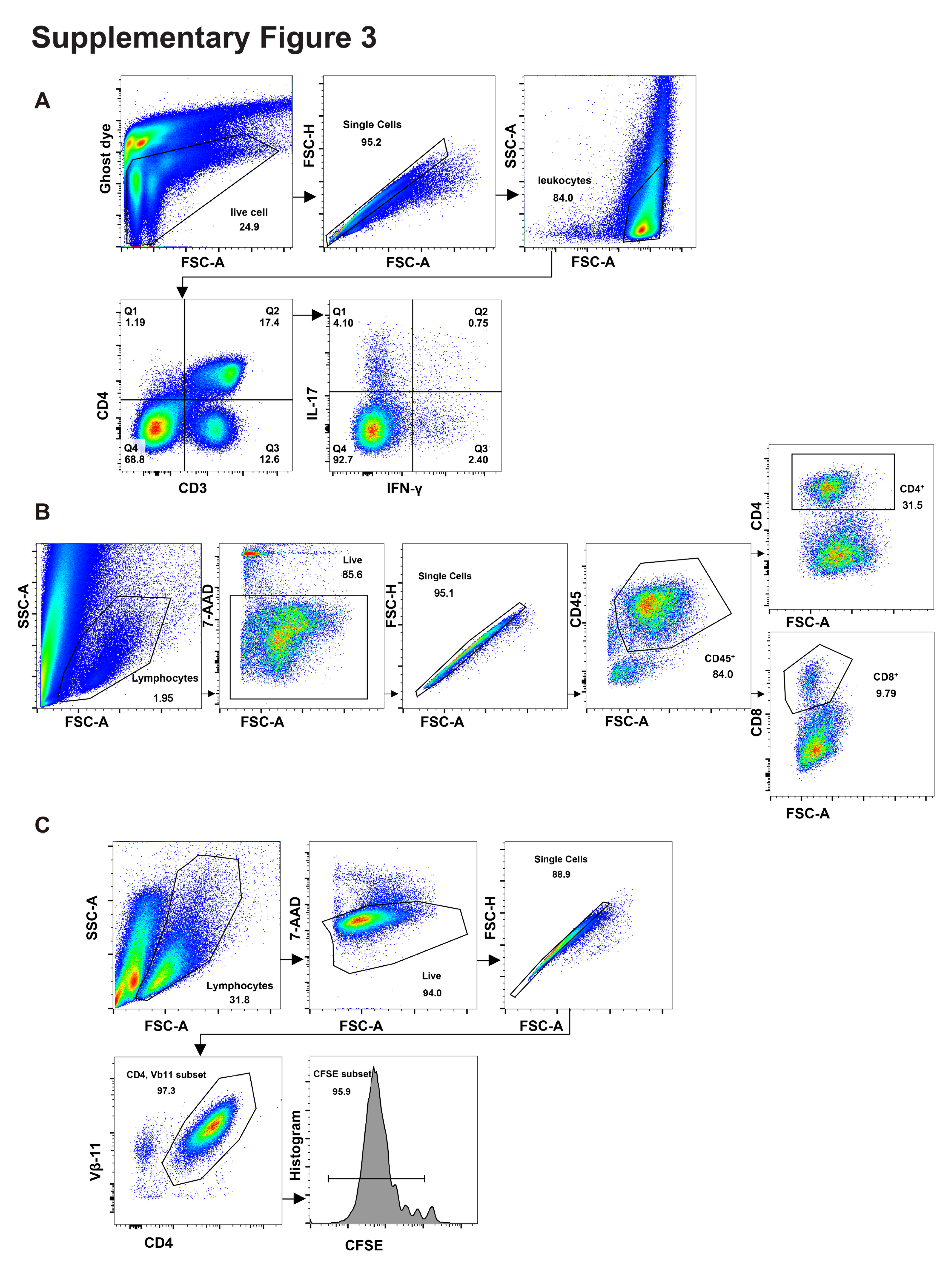
**Supplementary figure 3. Gating strategies.** (A) Gating strategy for cytokine response detection. (B) Gating strategy for the analysis of leukocytes in the spinal cord. (C) Gating strategy for the identification of CFSE population.

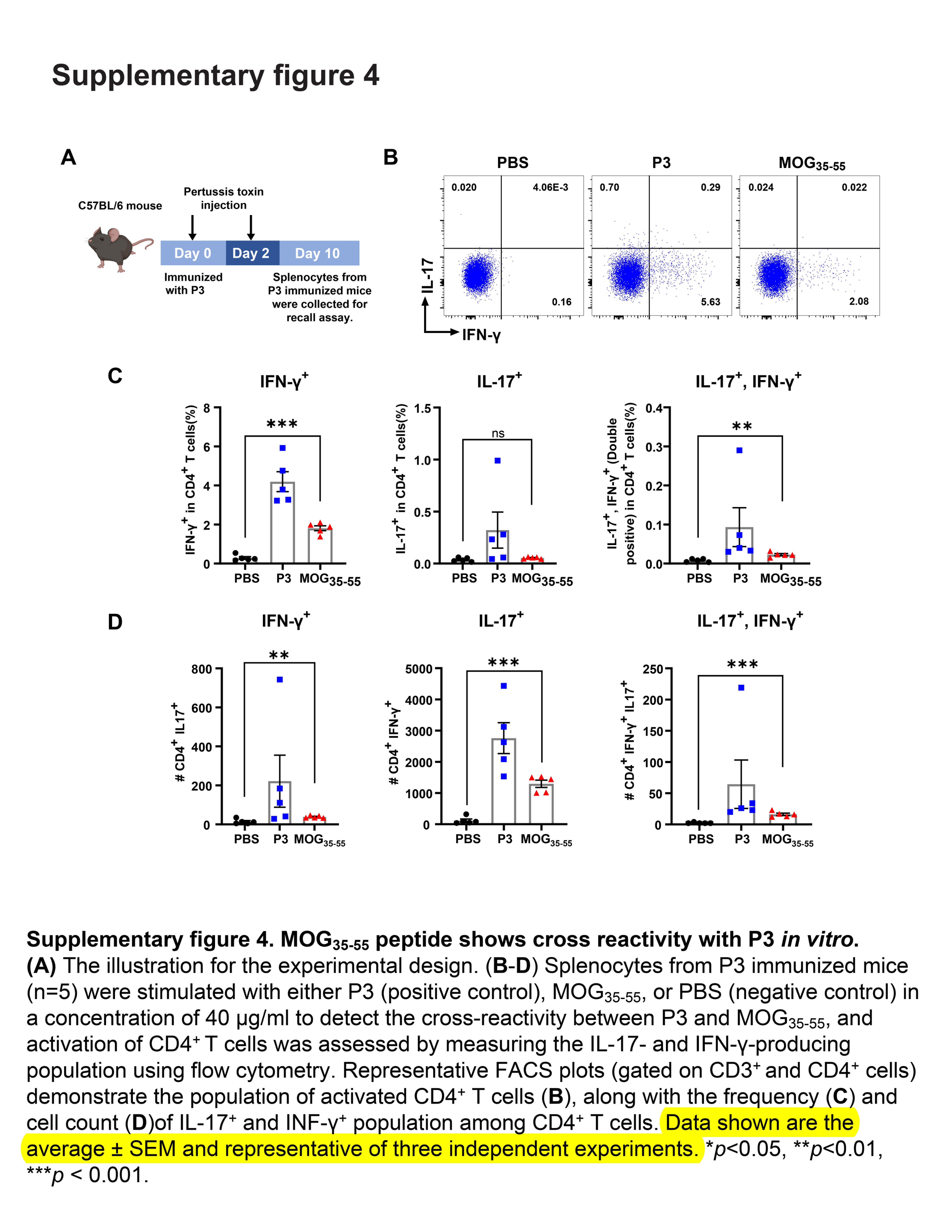
**Supplementary figure 4. MOG_35-55_ peptide shows cross reactivity with P3-specific T cells *in vitro*.** (A) Illustration of the experimental design. (B-D) Splenocytes from mice (n = 5) immunized with P3 were stimulated with either P3 (positive control), MOG_35-55_, or PBS (negative control), using a concentration of 20 μg/ml. The populations producing IL-17 and IFN-γ were assessed via flow cytometry. Representative FACS plots (B), the frequency (C), and cell count (D) of IL-17^+^ and IFN-γ^+^ cells within the CD4^+^ T cell population are presented. Results show one experiment that is representative of three independent experiments. The data shown are the average ± SEM. ***p*<0.01, ****p* < 0.001.

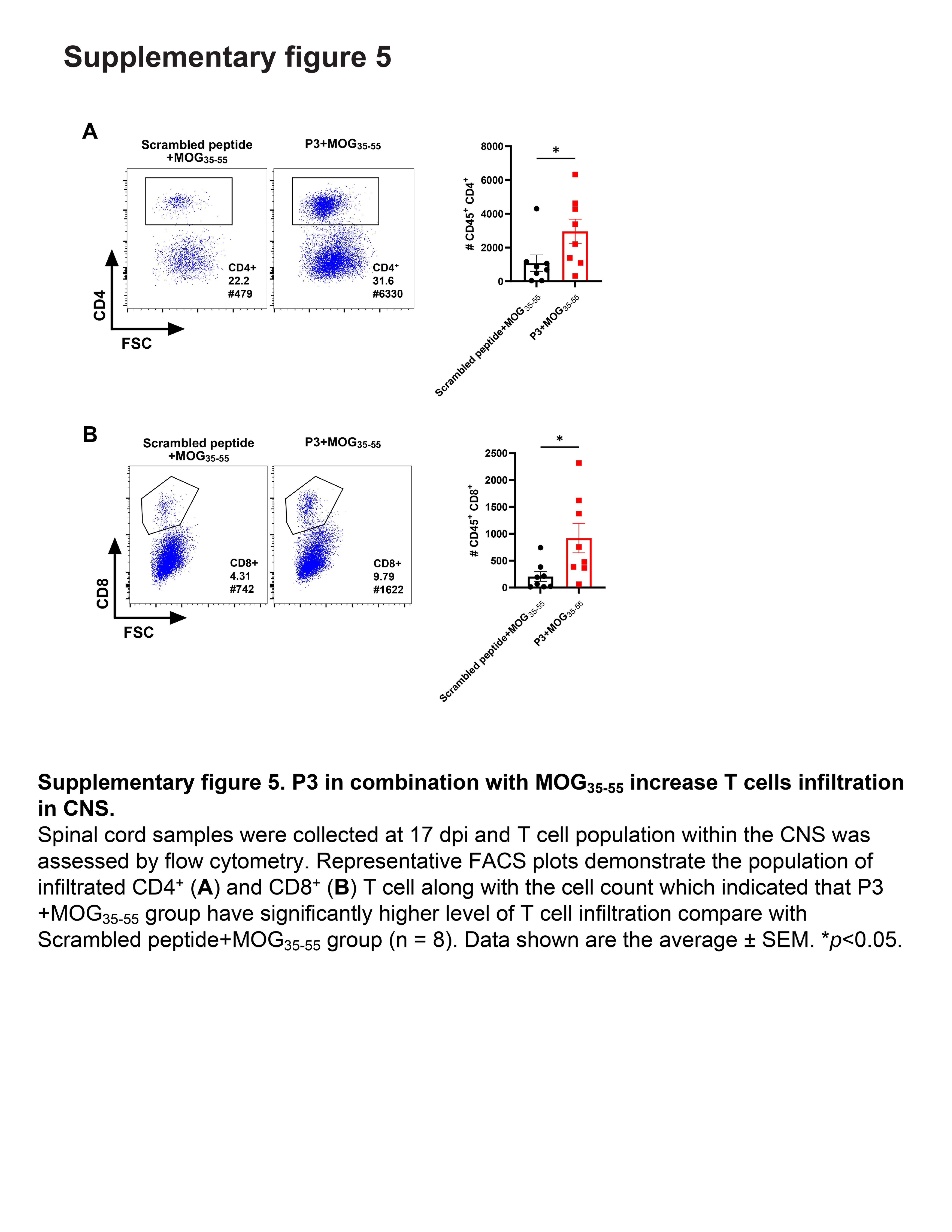
**Supplementary figure 5. P3 in combination with MOG_35-55_ increase T cells infiltration in CNS.** Spinal cord samples were collected at 17 dpi and T cell population within the CNS was assessed by flow cytometry. Representative FACS plots demonstrate the population of infiltrated CD4^+^ (A) and CD8^+^ (B) T cells. Data shown are the average ± SEM, n = 8 in each group. **p*<0.05.

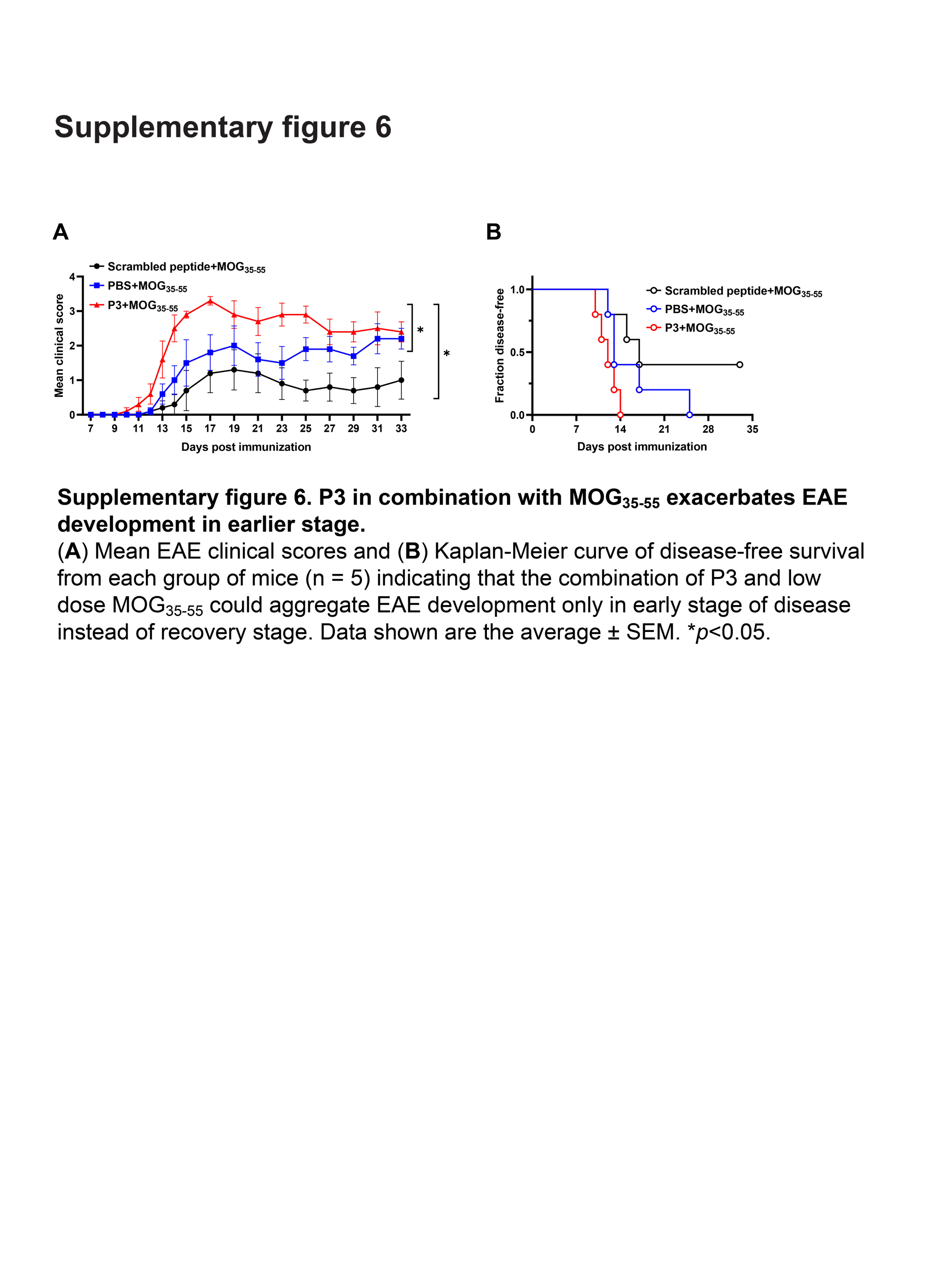
**Supplementary figure 6. P3 in combination with MOG_35-55_ exacerbates the development of EAE.** (A) Mean EAE clinical scores and (B) Kaplan-Meier curve of disease-free survival for each group of mice (n = 5) indicate that the combination of P3 and a low dose MOG_35-55_ enhances EAE progression during the early phase of the disease. Data are presented as the average ± SEM. **p* < 0.05.
